# The presenting HLA determines fidelity of SARS-CoV-2 spike protein epitope prediction

**DOI:** 10.1101/2024.07.24.604889

**Authors:** Charles R. Schutt, Deren Birol, Xiuyuan Lu, Sho Yamasaki

**Affiliations:** Laboratory of Molecular Immunology, Immunology Frontier Research Center, Osaka University, Suita, Japan; Department of Molecular Immunology, Research Institute for Microbial Diseases, Osaka University, Suita, Japan; Center for Infectious Disease Education and Research (CiDER), Osaka University, Suita, Japan

**Keywords:** epitope prediction, epitope presentation, SARS-CoV-2, spike protein, Helper T cells, epitopes, HLA

## Abstract

During the course of the COVID-19 pandemic, multiple studies used prediction methods to identify potential epitopes. While additional studies identified epitopes from convalescent and vaccinated subjects, few studies have compared the predicted to identified epitopes. Here we used three methods alone and in combination to predict helper T cell epitopes and compared the results to experimentally determined peptide binding. The correspondence between the results predicted from each method or combination and experimental results depends on the HLA being investigated. We were also able identify the prediction methods which lead to the most consistent results. Lastly, these observations were extended to more HLAs to predict epitopes which may be globally presented. All the predicted epitopes were previously identified as helper T cell epitopes. These results suggest predicting the binding to a larger number of HLAs may lead to higher fidelity identification of epitopes.

## Introduction

Several SARS-CoV-2 proteins are capable of eliciting strong humoral and cellular immune responses, among them is the spike glycoprotein (1–6). The spike protein consists of two subdomains, S1 (14–685) which contains the receptor binding domain (319–541), and S2 (686–1273) which mediates membrane fusion (7). In addition to facilitating viral entry (8), the spike protein contributes to pathology by multiple mechanisms including inducing thrombi, disrupting epithelial layers, and damaging the kidneys (9–11). Therefore, identifying the epitopes of the spike protein which generate the strongest anti-viral response and are conserved across variants is critical for optimizing the immune response against the SARS-CoV-2 spike protein.

Tools for predicting the binding of peptides to both HLA-I and HLA-II alleles are commonly available. Two frequently used methods are Consensus, whose prediction is the combined result from multiple top-performing tools, and NetMHCIIpan, which employs a machine learning model trained from datasets of peptides eluted from HLAs (EL) or binding affinity (BA) of the peptide for the HLA (12, 13). From the earliest days of the COVID-19 pandemic, researchers from around the globe began using these and other tools to predict epitopes which promote humoral and cellular immune responses (14–18). Since numerous studies have identified the epitopes recognized by the helper T cells of convalescent and vaccinated patients (1, 19–49), there is opportunity to test the performance of the different prediction methods alone or in combination.

In this study, we used prediction methods alone or combined to predict the binding of SARS-CoV-2 spike peptides to their binding to selected HLAs. Based on these results, we expanded our observations to identify the peptides which are predicted to be widely presented on the most HLAs by the greatest percentage of the global population.

## Results

Three different methods, IEDB recommended 2.22 (Consensus 2.22), NetMHCIIpan EL 4.0 and NetMHCIIpan BA 4.0 (12, 13) (Supplemental Figure 1), were used to predict the binding of SARS-CoV-2 spike proteins. We then confirmed the expression of each of five different HLAs (HLA-DRB1*15:01, HLA-DRB1*07:01, HLA-DRB1*04:05, HLA-DQA1*01:03/DQB1*06:01, and HLA-DPA1*02:02/DPB1*05:01) on separate cell lines (Supplemental Figure 2). These HLAs were chosen because they cover most of the Japanese and global population with minimal genetic linkage between the alleles (50, 51). The binding of biotinylated peptides to the HLA-expressing, but not non-expressing cells, was also confirmed (Supplemental Figures 2 and 3). The predicted and experimentally determined binding of each peptide to each HLA is shown in Supplemental Figure 1.

To determine the relationship between the predicted and experimental results, we first determined if there is a correlation between the predicted rank and the intensity of binding for each of the five HLAs tested (Figure 1). For HLA-DRB1*15:01 and HLA-DRB1*07:01, all three prediction methods significantly correlated. For HLA-DRB1*04:05, binding predicted by IEDB and NetMHCIIpan EL significantly correlated with experimental binding, while NetMHCIIpan BA did not. For HLA-DQA1*01:03/DQB1*06:01 and HLA-DPA1*02:02/DPB1*05:01 there was no correlation. Since the prediction methods yielded similar correlations for each HLA, the association between the predicted and experimentally derived results seems to depend more on the HLA than the prediction method used.

**Figure 1.**
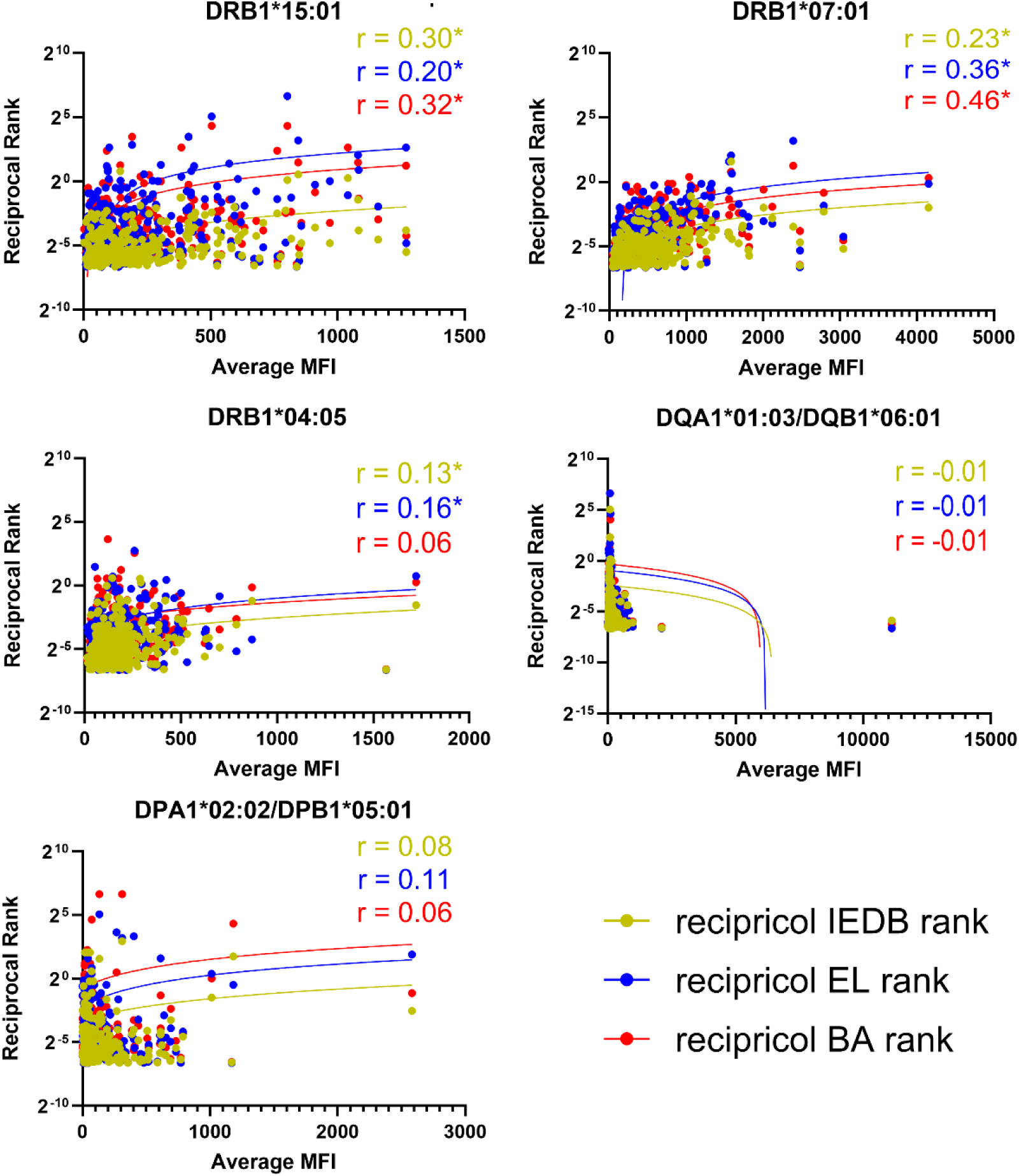
Correlation of predicted to experimental peptide binding on each HLA by prediction method For each HLA, the reciprocal rank was compared to the mean MFI for each peptide. Significance was determined by Spearman’s correlation coefficient. * p ≤ 0.05.

Peptides predicted to bind the HLA are expected to be found on the surface at higher levels than predicted non-binders. To test this, we compared the mean fluorescent intensity (MFI) of the predicted binders and non-binders as determined by each method alone or in combination. By all prediction methods or combinations, the predicted peptides had significantly higher MFI compared to predicted non-binders for HLA-DRB1*15:01 and HLA-DRB1*07:01 (Figure 2). Using NetMHCIIpan EL alone, NetMHCIIpan BA alone, or peptides predicated by any method, the predicted binding peptides to DRB1*04:05 had significantly higher MFI compared to predicted non-binders. For HLA-DQA1*01:03/DQB1*06:01 and HLA-DPA1*02:02/DPB1*05:01, there was no method or combination by which the predicted peptides had a higher MFI compared to non-binders. While the HLA of interest remains important for associating predicted peptides to surface binding, interestingly, refining predictions by using multiple methods did not lead to identification of more highly bound peptides.

**Figure 2.**
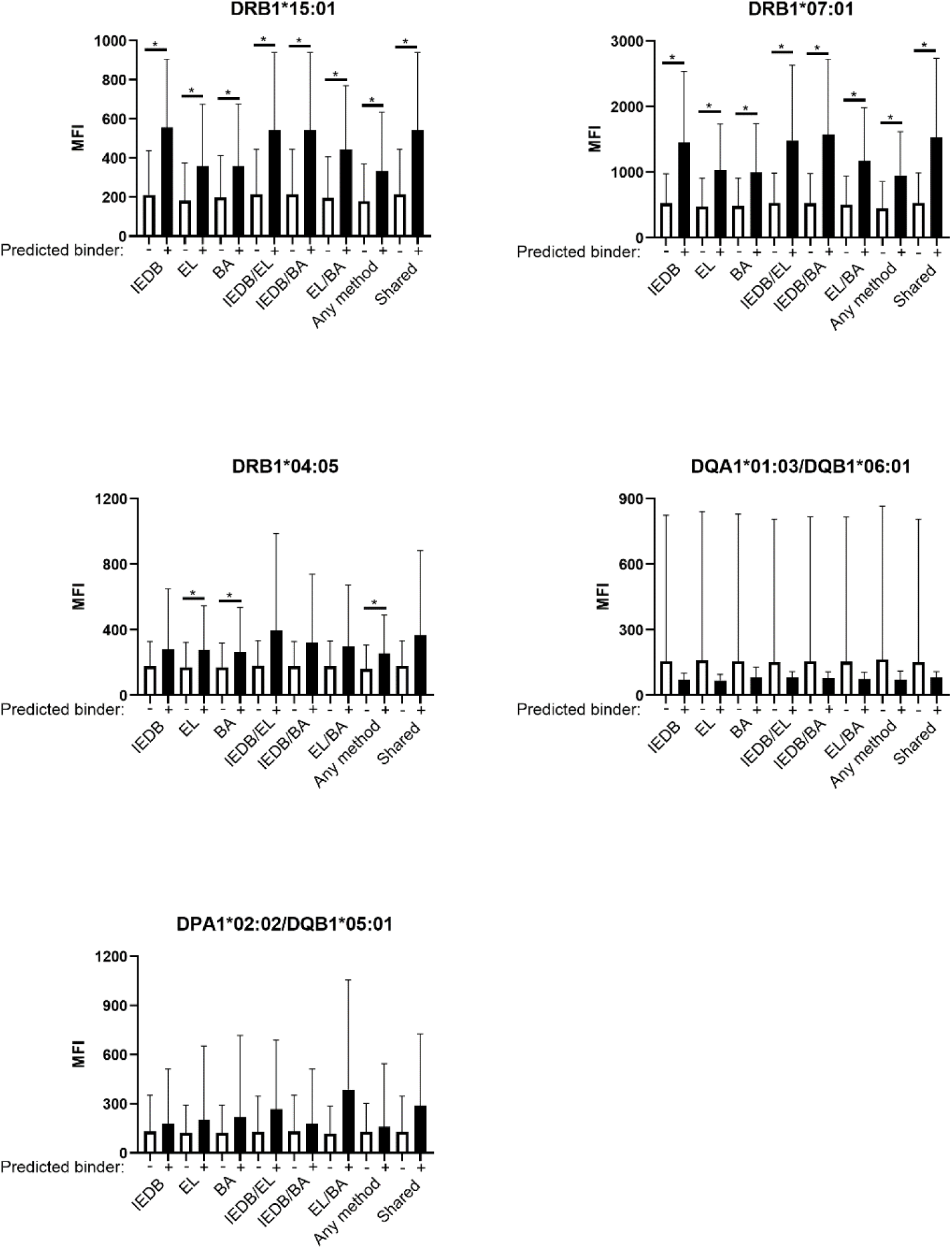
The MFI of predicted binding peptides for each HLA For each HLA, the MFI was compared for predicted binders and predicted non-binders as determined by each prediction method alone or in combination as indicated. Error bars are standard deviation. Significance was determined by Kruskal-Wallis test followed by Dunn’s Multiple comparisons test. * p ≤ 0.05.

We next sought to determine if the most highly bound peptides for each HLA (i.e. the peptides with the highest MFI) are concentrated among the predicted binding peptides. For HLA-DRB1*15:01 and HLA-DRB1*07:01, top bound peptides were significantly found among the predicted peptides, regardless of the prediction method or combination used (Table 1). For HLA-DRB1*04:05 and HLA-DPA1*02:02/DPB1*05:01, the number of bound peptides was significantly found among peptides predicted by NetMHCIIpan EL alone, NetMHCIIpan BA alone, these two methods in combination, and any peptide by any method. None of the top bound peptides to HLA-DQA1*01:03/DQB1*06:01 were predicted by any method or combination. As with Figure 2, the combination of methods did not improve the correspondence between the prediction and top bound peptides, as all significant associations involved the NetMHCIIpan methods.

**Table 1.** Contingency analysis of the number of top bound peptide with predicted peptides by each method.

|  | DRB1*15:01<sup>1-8</sup> | DRB1*07:01<sup>1-8</sup> | DRB1*04:05<sup>2,3,5,6,7</sup> | DQA1*01:03/DQB1*06:01 | DPA1*02:02/DPB1*05:01<sup>2,3,6,7</sup> |
| --- | --- | --- | --- | --- | --- |
| #predicted by IEDB | 12 | 12 | 22 | 23 | 19 |
| # of top 15 bound peptides predicted by IEDB | 5 | 4 | 2 | 0 | 3 |
| #predicted by EL | 76 | 46 | 41 | 34 | 41 |
| # of top 15 bound peptides predicted by EL | 10 | 6 | 5 | 0 | 5 |
| #predicted by BA | 50 | 46 | 47 | 25 | 33 |
| # of top 15 bound peptides predicted by BA | 7 | 6 | 7 | 0 | 5 |
| #predicted by IEDB/EL | 9 | 9 | 9 | 10 | 10 |
| # of top 15 bound peptides predicted by IEDB/EL | 4 | 3 | 1 | 0 | 2 |
| #predicted by IEDB/BA | 9 | 10 | 16 | 16 | 19 |
| # of top 15 bound peptides predicted by IEDB/BA | 4 | 3 | 3 | 0 | 2 |
| #predicted by EL/BA | 35 | 27 | 17 | 15 | 19 |
| # of top 15 bound peptides predicted by EL/BA | 4 | 3 | 1 | 0 | 2 |
| #predicted by any method | 95 | 66 | 77 | 50 | 56 |
| # of top 15 bound peptides predicted by any method | 11 | 8 | 8 | 0 | 4 |
| #predicted by all methods | 9 | 8 | 9 | 10 | 10 |
| # of top 15 presented peptides predicted by all methods | 4 | 3 | 1 | 0 | 2 |

For each HLA tested, the number of peptides predicted by all methods and combinations and the number of those peptides among the top 15 bound were listed. An association was significant if $p < 0.05$ using Fisher Exact Test. <sup>1</sup>IEDB (IEDB recommended 2.22), <sup>2</sup>EL (NetMHCIIpan EL 4.0), <sup>3</sup>BA (NetMHCIIpan BA 4.0), <sup>4</sup>IEDB/EL, <sup>5</sup>IEDB/BA, <sup>6</sup>EL/BA, <sup>7</sup>shared by all three methods, <sup>8</sup>any of the 3 methods. There were 315 total peptides.

Lastly, we sought to identify globally presented peptides by predicting binding to 206 HLA II alleles using the same methods. In order to identify the most peptides, we counted the number of HLAs predicted to bind each peptide by any of the prediction methods and 30 peptides (33, 37, 41, 53, 57, 229, 233, 237, 309, 313, 337, 341, 345, 509, 689, 761, 765, 777, 797, 853, 893, 897, 917, 1005, 1009, 1013, 1017, 1057, and 1061) were predicted to bind to more than half of the HLAs (Figure 3). Between 18-97% of the world may express an HLA-DRB1 allele predicted to present these peptides. To determine if the predicted peptides are actual epitopes, we examined the literature of the identified spike protein helper T cell epitopes. All 30 of the predicted peptides were found in at least one source with 3 peptides (233, 449, and 1009) commonly identified in the literature. These peptides are predicted to bind to HLA-DRB1s expressed by 82, 68, and 34% of the global population respectively. Using this broad approach, the predicted peptides were previously identified as helper T cell epitopes from convalescent or vaccinated individuals.

**Figure 3.**
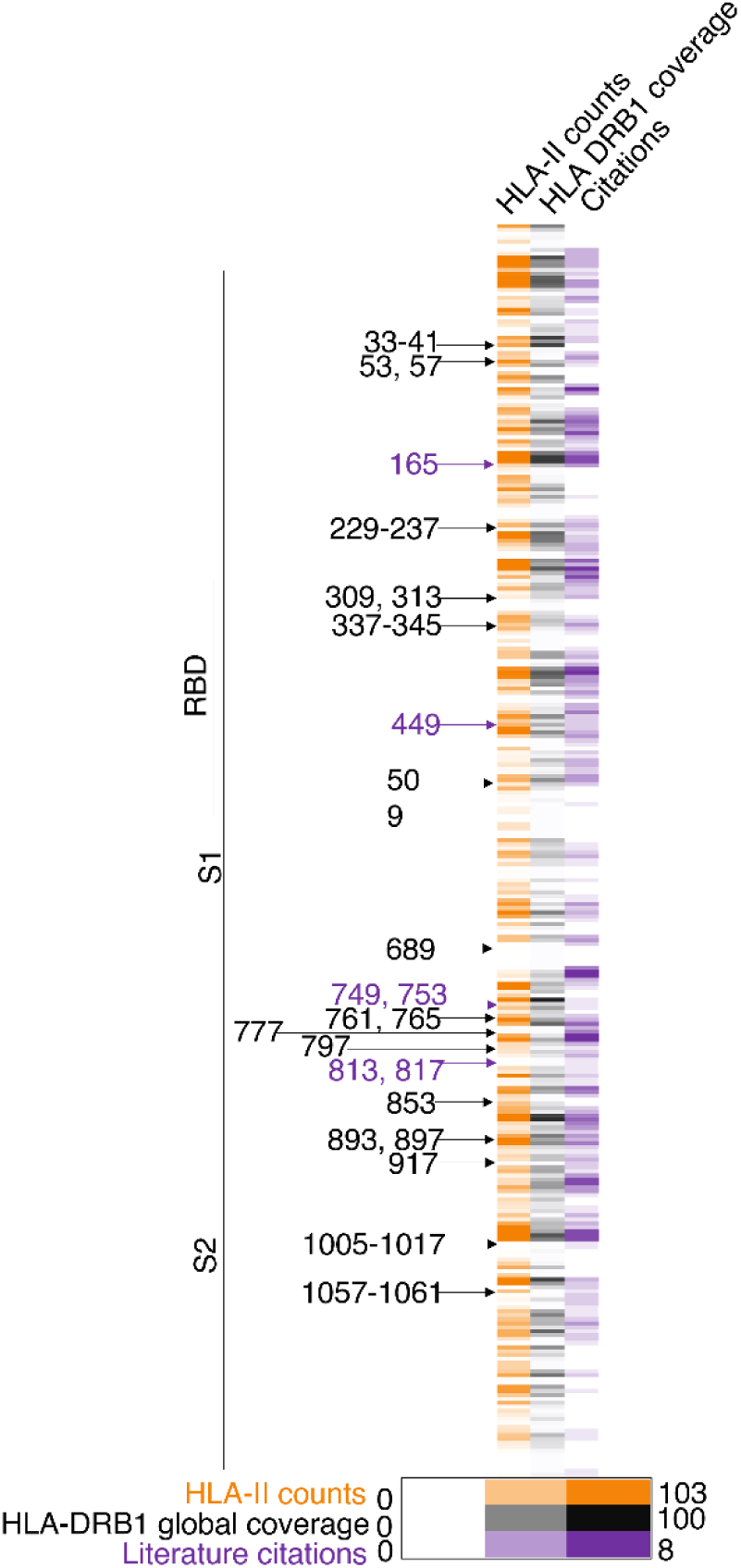
Predicting the presentation of SARS-CoV-2 spike protein on a wider number of HLAs A peptide is considered to bind to each HLA if one of the 12-, 13-, 14-, or 15-mers for each peptide is predicted to bind by at least one of the methods. All unique HLAs predicted to bind a peptide were counted (orange). The percentage of the world population which expresses HLA-DRB1s predicted to bind each peptide (black). The number of citations which identify each peptide as a T cell epitope was determined (purple). The black arrows indicate peptides predicted to bind more than half of the HLAs. The purple arrows indicate peptides found commonly in the literature. The location of the S1 region, receptor binding domain (RBD) and S2 region are indicated on the left.

## Discussion

In this study we co-cultured biotinylated 15-mers with HLA-expressing cells to measure peptide binding. However, we did not establish a level of binding in this system which is needed to identify epitopes. As highlighted by the red arrow/arrowhead pairs in Supplemental Figure 1, many of the most highly bound peptides were also previously identified epitopes. This suggests that the peptides which are highly bound in this system may be epitopes, even if not previously predicted or identified. Many of the remaining peptides were predicted to be epitopes by at least one of the methods and bound to the HLA to varying degrees. However, there were several identified epitopes (297, 301, 849, and 853 for HLA-DRB1*07:01, 481 and 485 for HLA-DRB1*04:05, and 993 for HLA-DQA1*01:03/DQB1*06:01) (25, 36, 48), which are not among the top bound peptides nor were they predicted to bind. Therefore, our system was able to identify previously published helper T cell epitopes, but more work is needed to further refine this method.

Within our system, there are discrepancies between the predictions and the binding of peptides. The peptides which are predicted to be presented on a large number of HLAs are generally not found among the most highly bound peptides in our system. Furthermore, some of the most highly bound peptides to the five HLAs in this study (Figure 1) were not previously predicted to be an epitope (for example 557, 713, 717, 733, and 1245). Some peptides (733 and 1245) have a higher background (Supplemental Figure 3), so it is possible that these peptides are nonspecifically binding to the cell and/or to the HLA. Other peptides (557, 713, and 717) do not have such high background and may be more likely to be presented. This aligns with the literature as peptides 557, 713, and 717 (24, 27, 35, 48) have been identified as epitopes while 733 and 1245 were not (Figure 3).

Among the 30 peptides which were predicted to be broadly presented, many contain few or no known mutations in the recent or currently circulating variants of interest, such as XBB.1.5, XBB1.16, EG.5, BA.2.75, BA.2.86, JN.1. The lack of mutations in these peptides suggest that they are located in regions conserved across variants. However, recent and currently circulating variants of interest contain mutations in peptides including 37, 41, 57, 237, 337, 341, 345, 449, and 761. While some of these mutations are predicted to increase the number of presenting HLAs (Supplemental Figure 4), several mutations (such as those in peptides 337 and 494), are predicted to bind to noticeably fewer HLAs. Interestingly, these peptides are located in the receptor binding domain (7). This observation is in line with humoral data in which BA.2.86 and JN.1 are less recognized by antibodies generated by prior infection or vaccination (52–55). However, these mutations do not appear sufficient for T cell immunity escape (56, 57). Therefore, it is possible that cellular immunity obtained by infection and/or vaccination may continue even if the humoral response diminishes.

In this study, we utilized three different prediction methods to identify potential helper T cell epitopes. These different methods utilize different strategies to make predictions as well as being trained using different data sets (12, 13). However, the three HLA-DRB1 alleles were included in all of the training datasets, but the HLA-DQ and HLA-DP alleles were not, which may explain the differences in the degree of association with these alleles. Of the methods used here, NetMHCIIpan EL and BA gave the most consistent results across the HLAs tested (Figures 1, 2, and Table 1), suggesting that expanding the datasets of eluted peptides and binding affinities may be key to improving the accuracy of predictions. This may also suggest the machine learning approach used by NetMHCIIpan may provide more consistent results than the matrix-based approach used by Consensus. Furthermore, the correspondence between predicted and experimental binding was variable when examining individual HLAs, but higher when predicting binding a larger number of HLAs. This suggests that at this time, these prediction methods may be better at identifying epitopes generally than for specific HLAs.

## Methods

### Prediction methods

The Wuhan SARS CoV-2 spike protein sequence (Uniprot P0DTC2) was divided into 15-mer peptides with 11 amino acid overlap. Each peptide is numbered by the position of the first amino acid in the peptide.

All predictions were made using Immune Epitope Database and Tools (IEDB) prediction methods: IEDB recommended 2.22 (Consensus 2.22), NetMHCIIpan EL 4.0 and NetMHCIIpan BA 4.0. For each peptide, the binding of 12-, 13-, 14-, and 15-mer of each peptide was predicted with all three methods. A peptide was considered to bind if the reciprocal rank for any of the 12-15-mers was 0.2 or greater. The highest reciprocal ranked 12-15-mer for each peptide was presented in Figure 1 and supplemental Figure 1.

We tested the binding to the following 206 HLA-II alleles:

HLA-DRB1*01:01, HLA-DRB1*01:02, HLA-DRB1*03:01, HLA-DRB1*04:01, HLA-DRB1*04:02, HLA-DRB1*04:03, HLA-DRB1*04:04, HLA-DRB1*04:05, HLA-DRB1*04:06, HLA-DRB1*04:07, HLA-DRB1*04:08, HLA-DRB1*04:10, HLA-DRB1*04:11, HLA-DRB1*04:57, HLA-DRB1*07:01, HLA-DRB1*08:01, HLA-DRB1*08:02, HLA-DRB1*08:03, HLA-DRB1*08:04, HLA-DRB1*08:09, HLA-DRB1*08:23, HLA-DRB1*09:01, HLA-DRB1*10:01, HLA-DRB1*11:01, HLA-DRB1*11:04, HLA-DRB1*11:06, HLA-DRB1*11:08, HLA-DRB1*12:01, HLA-DRB1*12:02, HLA-DRB1*12:05, HLA-DRB1*13:01, HLA-DRB1*13:02, HLA-DRB1*13:03, HLA-DRB1*13:07, HLA-DRB1*14:01, HLA-DRB1*14:02, HLA-DRB1*14:03, HLA-DRB1*14:04, HLA-DRB1*14:05, HLA-DRB1*14:06, HLA-DRB1*14:07, HLA-DRB1*14:12, HLA-DRB1*14:29, HLA-DRB1*14:45, HLA-DRB1*14:54, HLA-DRB1*15:01, HLA-DRB1*15:02, HLA-DRB1*15:03, HLA-DRB1*15:04, HLA-DRB1*15:06, HLA-DRB1*15:11, HLA-DRB1*16:01, HLA-DRB1*16:02, HLA-DRB3*03:01, HLA-DRB3*01:01, HLA-DRB3*02:02, HLA-DRB4*01:01, HLA-DRB4*01:03, HLA-DRB4*01:04, HLA-DRB5*01:01, HLA-DRB5*01:02, HLA-DRB5*02:02, HLA-DQA1*05:01/DQB1*02:01, HLA-DQA1*06:01/DQB1*02:01, HLA-DQA1*02:01/DQB1*02:01, HLA-DQA1*02:01/DQB1*02:02, HLA-DQA1*03:03/DQB1*02:02, HLA-DQA1*03:03/DQB1*02:03, HLA-DQA1*03:02/DQB1*03:01, HLA-DQA1*05:01/DQB1*03:01, HLA-DQA1*05:05/DQB1*03:01, HLA-DQA1*05:03/DQB1*03:01, HLA-DQA1*06:01/DQB1*03:01, HLA-DQA1*03:03/DQB1*03:01, HLA-DQA1*05:08/DQB1*03:01, HLA-DQA1*05:06/DQB1*03:01, HLA-DQA1*01:03/DQB1*03:01, HLA-DQA1*03:01/DQB1*03:01, HLA-DQA1*03:02/DQB1*03:02, HLA-DQA1*03:01/DQB1*03:02, HLA-DQA1*04:01/DQB1*03:02, HLA-DQA1*03:03/DQB1*03:02, HLA-DQA1*05:05/DQB1*03:02, HLA-DQA1*05:03/DQB1*03:02, HLA-DQA1*05:08/DQB1*03:02, HLA-DQA1*02:01/DQB1*03:03, HLA-DQA1*03:01/DQB1*03:03, HLA-DQA1*03:02/DQB1*03:03, HLA-DQA1*03:03/DQB1*03:03, HLA-DQA1*01:01/DQB1*03:03, HLA-DQA1*05:05/DQB1*03:03, HLA-DQA1*06:01/DQB1*03:03, HLA-DQA1*05:03/DQB1*03:19, HLA-DQA1*05:05/DQB1*03:19, HLA-DQA1*03:03/DQB1*04:01, HLA-DQA1*03:01/DQB1*04:01, HLA-DQA1*04:01/DQB1*04:01, HLA-DQA1*05:01/DQB1*04:01, HLA-DQA1*02:01/DQB1*04:02, HLA-DQA1*03:03/DQB1*04:02, HLA-DQA1*04:01/DQB1*04:02, HLA-DQA1*03:01/DQB1*04:02, HLA-DQA1*05:01/DQB1*04:02, HLA-DQA1*06:01/DQB1*04:02, HLA-DQA1*01:01/DQB1*05:01, HLA-DQA1*01:02/DQB1*05:01, HLA-DQA1*01:03/DQB1*05:01, HLA-DQA1*01:04/DQB1*05:01, HLA-DQA1*01:05/DQB1*05:01, HLA-DQA1*02:01/DQB1*05:01, HLA-DQA1*01:03/DQB1*05:03, HLA-DQA1*01:04/DQB1*05:03, HLA-DQA1*03:03/DQB1*05:03, HLA-DQA1*01:04/DQB1*05:02, HLA-DQA1*01:01/DQB1*05:02, HLA-DQA1*01:02/DQB1*05:02, HLA-DQA1*01:01/DQB1*05:03, HLA-DQA1*01:03/DQB1*06:01, HLA-DQA1*01:02/DQB1*06:01, HLA-DQA1*03:01/DQB1*06:01, HLA-DQA1*05:01/DQB1*06:01, HLA-DQA1*05:03/DQB1*06:01, HLA-DQA1*01:02/DQB1*06:02, HLA-DQA1*01:03/DQB1*06:02, HLA-DQA1*05:03/DQB1*06:02, HLA-DQA1*01:03/DQB1*06:03, HLA-DQA1*06:01/DQB1*06:03, HLA-DQA1*01:02/DQB1*06:04, HLA-DQA1*01:03/DQB1*06:04, HLA-DQA1*03:03/DQB1*06:04, HLA-DQA1*01:02/DQB1*06:09, HLA-DQA1*01:03/DQB1*06:09, HLA-DPA1*02:02/DPB1*01:01, HLA-DPA1*02:01/DPB1*01:01, HLA-DPA1*01:03/DPB1*02:01, HLA-DPA1*02:02/DPB1*02:01, HLA-DPA1*02:01/DPB1*02:01, HLA-DPA1*02:02/DPB1*02:02, HLA-DPA1*01:03/DPB1*02:02, HLA-DPA1*02:02/DPB1*03:01, HLA-DPA1*01:03/DPB1*03:01, HLA-DPA1*01:03/DPB1*04:01, HLA-DPA1*02:01/DPB1*04:01, HLA-DPA1*02:02/DPB1*04:01, HLA-DPA1*01:03/DPB1*04:02, HLA-DPA1*02:02/DPB1*04:02, HLA-DPA1*04:01/DPB1*04:02, HLA-DPA1*02:02/DPB1*05:01, HLA-DPA1*02:01/DPB1*05:01, HLA-DPA1*01:03/DPB1*05:01, HLA-DPA1*01:03/DPB1*06:01, HLA-DPA1*02:01/DPB1*09:01, HLA-DPA1*02:01/DPB1*10:01, HLA-DPA1*02:02/DPB1*10:01, HLA-DPA1*02:02/DPB1*11:01, HLA-DPA1*02:01/DPB1*104:01, HLA-DPA1*02:02/DPB1*104:01, HLA-DPA1*01:03/DPB1*105:01, HLA-DPA1*02:01/DPB1*105:01, HLA-DPA1*02:01/DPB1*13:01, HLA-DPA1*04:01/DPB1*13:01, HLA-DPA1*04:01/DPB1*14:01, HLA-DPA1*02:01/DPB1*14:01, HLA-DPA1*02:02/DPB1*14:01, HLA-DPA1*02:01/DPB1*17:01, HLA-DPA1*02:01/DPB1*19:01, HLA-DPA1*02:02/DPB1*19:01, HLA-DPA1*01:03/DPB1*20:01, HLA-DPA1*01:03/DPB1*21:01, HLA-DPA1*01:03/DPB1*25:01, HLA-DPA1*02:02/DPB1*27:01, HLA-DPA1*04:01/DPB1*28:01, HLA-DPA1*02:02/DPB1*29:01, HLA-DPA1*02:02/DPB1*31:01, HLA-DPA1*01:03/DPB1*36:01, HLA-DPA1*02:02/DPB1*38:01, HLA-DPA1*01:03/DPB1*41:01, HLA-DPA1*01:03/DPB1*18:01, HLA-DPA1*01:03/DPB1*77:01, HLA-DPA1*01:03/DPB1*14:01, HLA-DPA1*01:03/DPB1*33:01, HLA-DPA1*01:03/DPB1*11:01, HLA-DPA1*01:03/DPB1*104:01, HLA-DPA1*01:03/DPB1*13:01, HLA-DPA1*01:03/DPB1*24:01, HLA-DPA1*01:03/DPB1*23:01, HLA-DPA1*01:03/DPB1*15:01, HLA-DPA1*01:03/DPB1*30:01, HLA-DPA1*01:03/DPB1*49:01, HLA-DPA1*01:03/DPB1*19:01, HLA-DPA1*01:03/DPB1*16:01, HLA-DPA1*01:03/DPB1*17:01, HLA-DPA1*02:01/DPB1*18:01, HLA-DPA1*02:01/DPB1*26:01, HLA-DPA1*02:01/DPB1*76:01, HLA-DPA1*02:01/DPB1*70:01, HLA-DPA1*03:01/DPB1*04:02, HLA-DPA1*03:01/DPB1*105:01, HLA-DPA1*03:01/DPB1*01:01, HLA-DPA1*03:01/DPB1*04:01, HLA-DPA1*03:01/DPB1*60:01, HLA-DPA1*03:01/DPB1*55:01, HLA-DPA1*03:01/DPB1*40:01, HLA-DPA1*03:01/DPB1*03:01, HLA-DPA1*03:01/DPB1*02:01, and HLA-DPA1*03:01/DPB1*19:01.

### FACS

A mouse T cell hybridoma cell line was transfected as previously described (58). Briefly, the HLA α and β cDNA sequences were synthesized and cloned into retroviral vectors which were transfected into Phoenix packaging cells using PEI MAX (Polysciences). Supernatant containing retroviruses was used to infect mouse T cell hybridoma cells. HLA expression was confirmed by flow cytometry.

All peptides were synthesized with a GSGSGS tag added to the N terminus and biotinylated. To test the binding of each peptide, 3.5 μg of biotinylated peptide was added per well of a 96 well U-bottom plate along with 2.5 × 10^4^ HLA null and 2.5 × 10^4^ HLA-expressing T cell hybridoma cells. After at least 18 hours of culture at 37°C for 5% CO_2_, cells were stained with streptavidin-conjugated R-phycoerythrin (SA-PE) (Biolegend cat# 405204) and HLA-DR/DP/DQ conjugated to APC (Biolegend clone Tϋ39 cat#361714) at 4°C for 45 min. After washing, cells were resuspended in 0.5x propidium iodine (Sigma P4170-100mg) prior to performing flow cytometry using the Gallios (Beckman Coulter). Background adjusted median fluorescent intensity (MFI) was calculated by subtracting the intensity from HLA-null cells from the HLA-expressing HLA cells in the same well. Experiments were repeated three times for each HLA and the average of all three replicates are presented in Figure 1, 2 and Supplemental Figure 1.

### Statistics

Correlation was determined by Spearman’s correlation coefficient. Significance was determined if the p value ≤ 0.05. Comparisons between groups was done with Kruskal-Wallis test followed by Dunn’s Multiple comparisons test. A p value ≤ 0.05 is considered significant. Contingency analysis was performed using Fisher’s exact test. A p value ≤ 0.05 is considered significant. All statistics were performed with GraphPad Prism version 10.2.

## Supplemental Figures

**Supplemental Figure 1.**
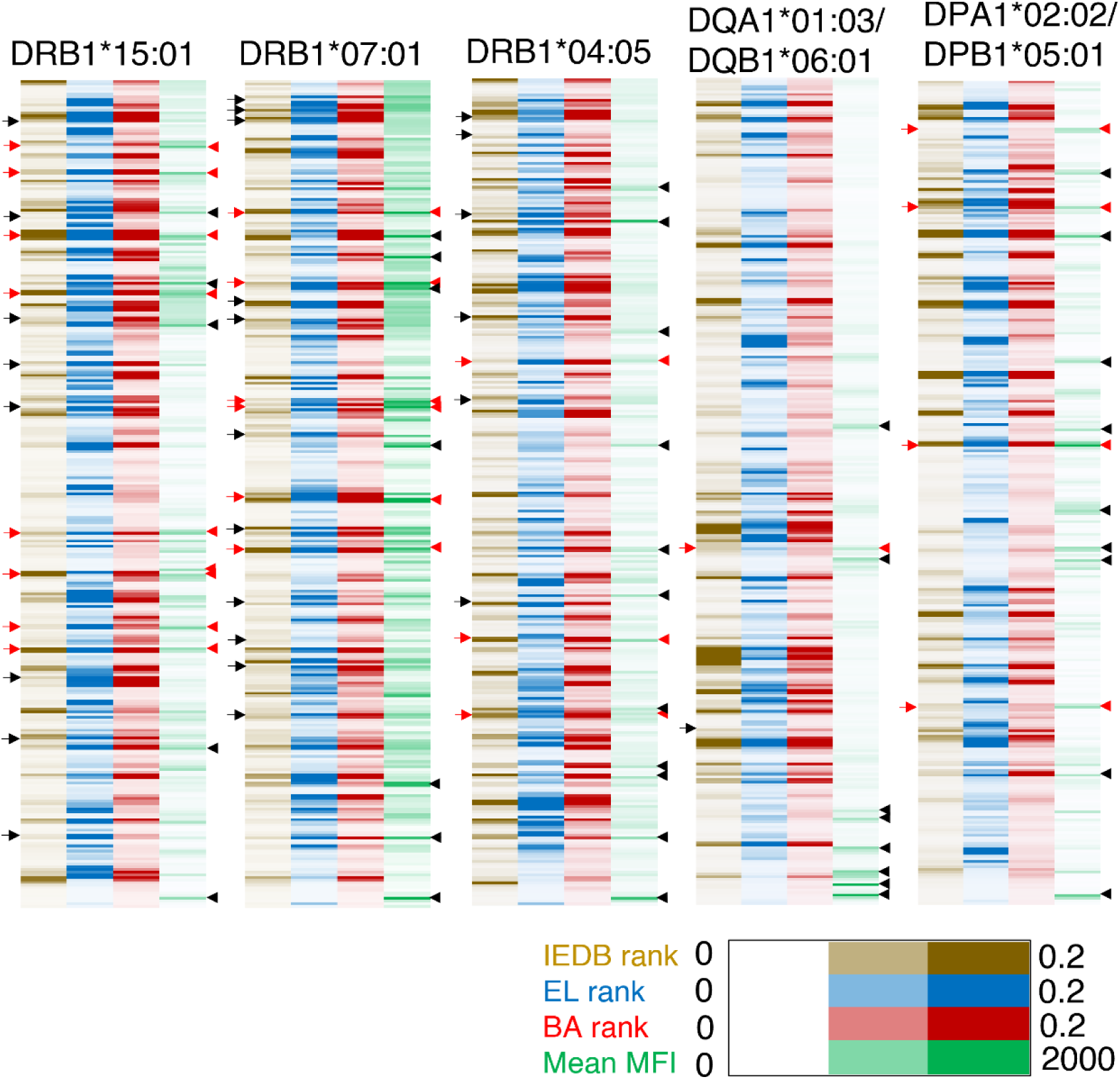
Comparison of the predicted reciprocal rank of SARS-CoV-2 spike peptides with the binding to each HLA For each of the HLAs, the reciprocal rank of the highest ranked 12-, 13-, 14- and 15-mer for each peptide predicted by IEDB recommended 2.22 (yellow), NetMHCIIpan EL 4.0 (blue), NetMHCIIpan BA 4.0 (red) along with the mean background-corrected MFI of T cell hybridoma cells (green) expressing the HLA of interest cultured with the biotinylated peptide of interest are presented. The arrows on the left are highlighting the peptides which are epitopes identified in the literature. The arrow heads on the right are highlighting the 15 peptides with the highest MFI. The red arrows and arrowheads indicate shared peptides. For HLA-DRB1*15:01, the arrows on the left are at peptides 57/61/65, 97/101/105, 141, 205/209, 233/237/241, 321/325, 361/365, 425/429/433/437/441/445, 497/501, 689/693/697, 745/749/753/757/761, 833, 865/869/873, 905/909/913/917/921, 1001/1005/1009, and 1153 and the arrow heads on the right are peptides 101, 141, 201, 237, 309, 321/325, 373, 689, 745, 753, 833, 865, 1017, and 1245. For HLA-DRB1*07:01, the arrows on the left correspond 25/29/33, 41/45/49, 57/61/65, 193/197/201/205/209, 297/301/305/309/313, 337, 361/365, 489/493/497/501, 537/541, 625/629/633/637/641, 673/677/681/685/689/693/697, 713/717/721, 793/797, 849/853, 889/893/897, and 965/969 and the arrowheads on the right correspond to peptides 201, 237, 269, 309, 317, 489, 497, 557, 637/641, 713, 1069/1073, 1153, and 1245. For HLA-DRRB1*04:05, the arrows on the left correspond to peptides 57/61, 85/89, 205/209, 361/365, 429/433, 481/485/489/493, 793/797, 849/853, and 965/969 and the arrowheads on the right correspond to 165/169, 217, 385, 429, 557, 717, 785, 853, 957, 965, 1045, 1061, 1153, 1245. For HLA-DQA*01:03/DQB1*06:01, the arrows on the left correspond to peptides 717/721 and 993 and the arrowheads on the right correspond to peptides 529/533, 717, 733/737, 1117, 1129, 1173/1177, 1209/1213/1217, 1229, and 1245/1249. For HLA-DPA1*02:02/DPB1*05:01, the arrows on the left correspond to peptides 73/77, 193, 553/557/561, and 953/957/961 and the arrowheads on the right correspond to peptides 73, 141, 193, 237, 429, 533, 553/557, 657, 713/717, 733, 957, 1061, and 1245.

**Supplemental Figure 2.**
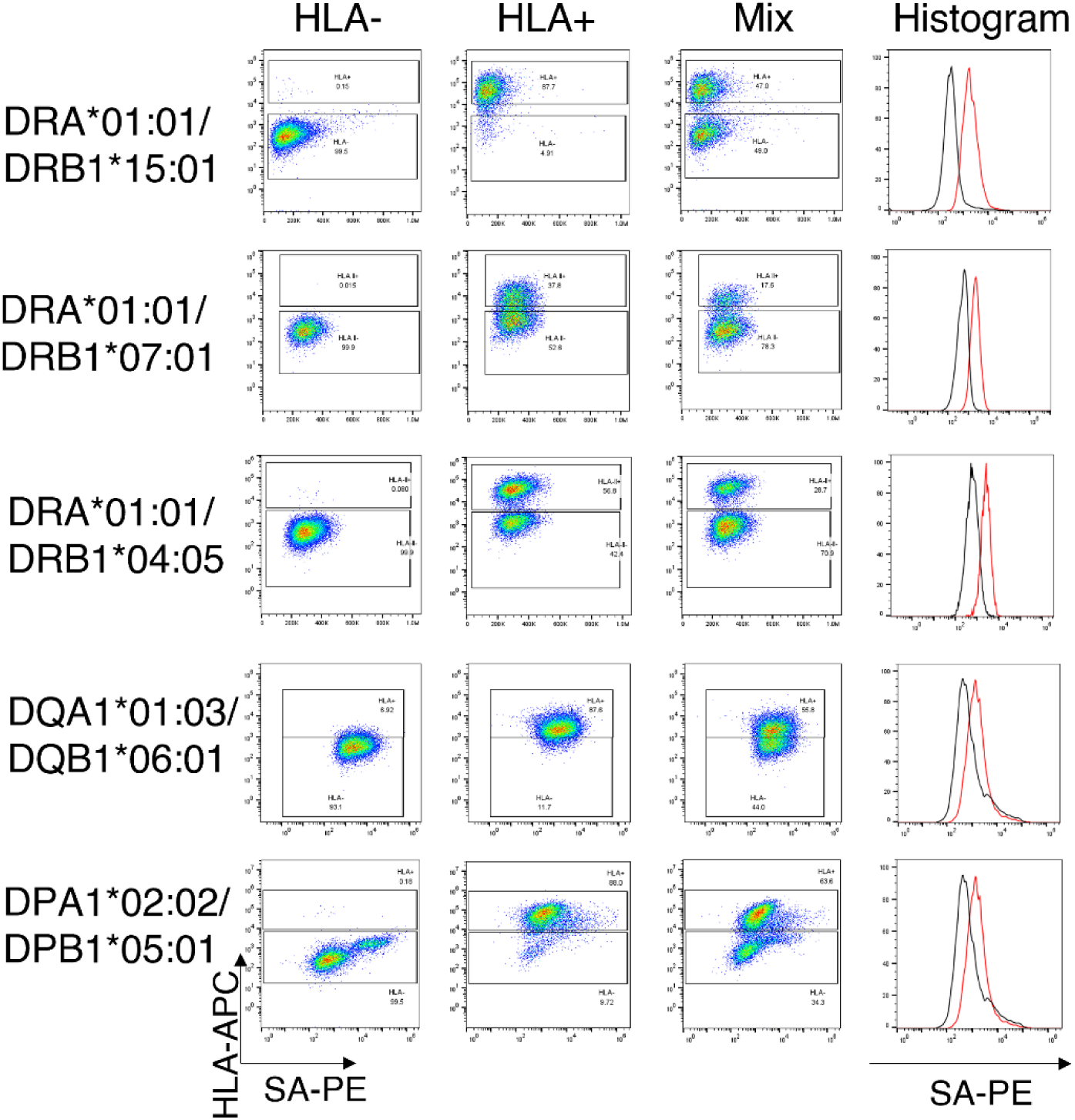
Confirming HLA expression For all five HLAs tested, representative images of HLA null (HLA-), HLA-expressing (HLA+), mixture of both cells, and histograms HLA-(black) and HLA+ (red) are shown. All cells were cultured with one of the top bound peptides for the indicted HLA (HLA-DRB1*15:01 - peptide 201, HLA-DRB1*07:01 - peptide 237, HLA-DRB1*04:05 -peptide 217, HLA-DQA1*01:03/DQB1*06:01 - peptide 733, HLA-DPA1*02:02/DPB1*05:01 - peptide 237).

**Supplemental Figure 3.**
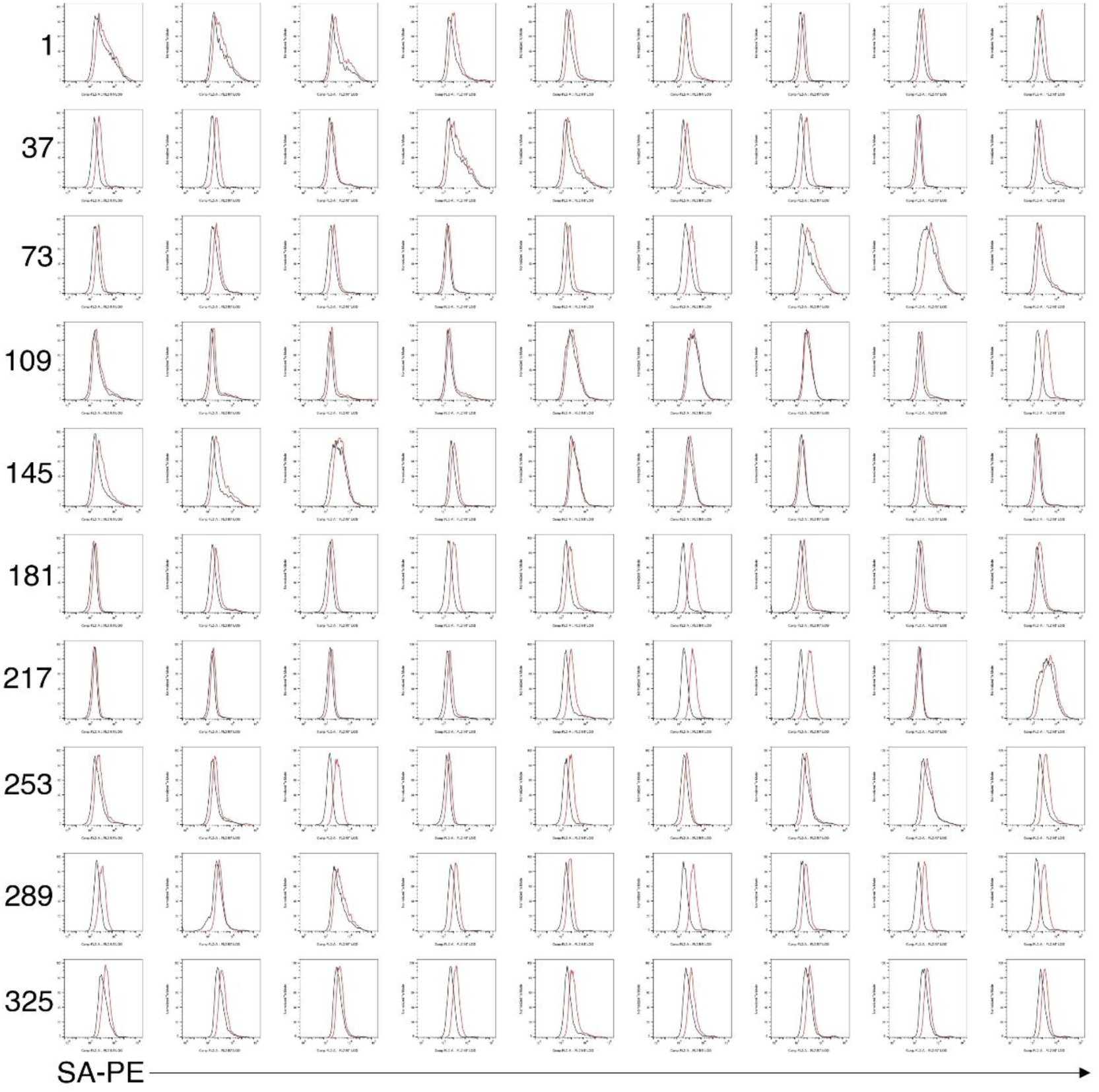
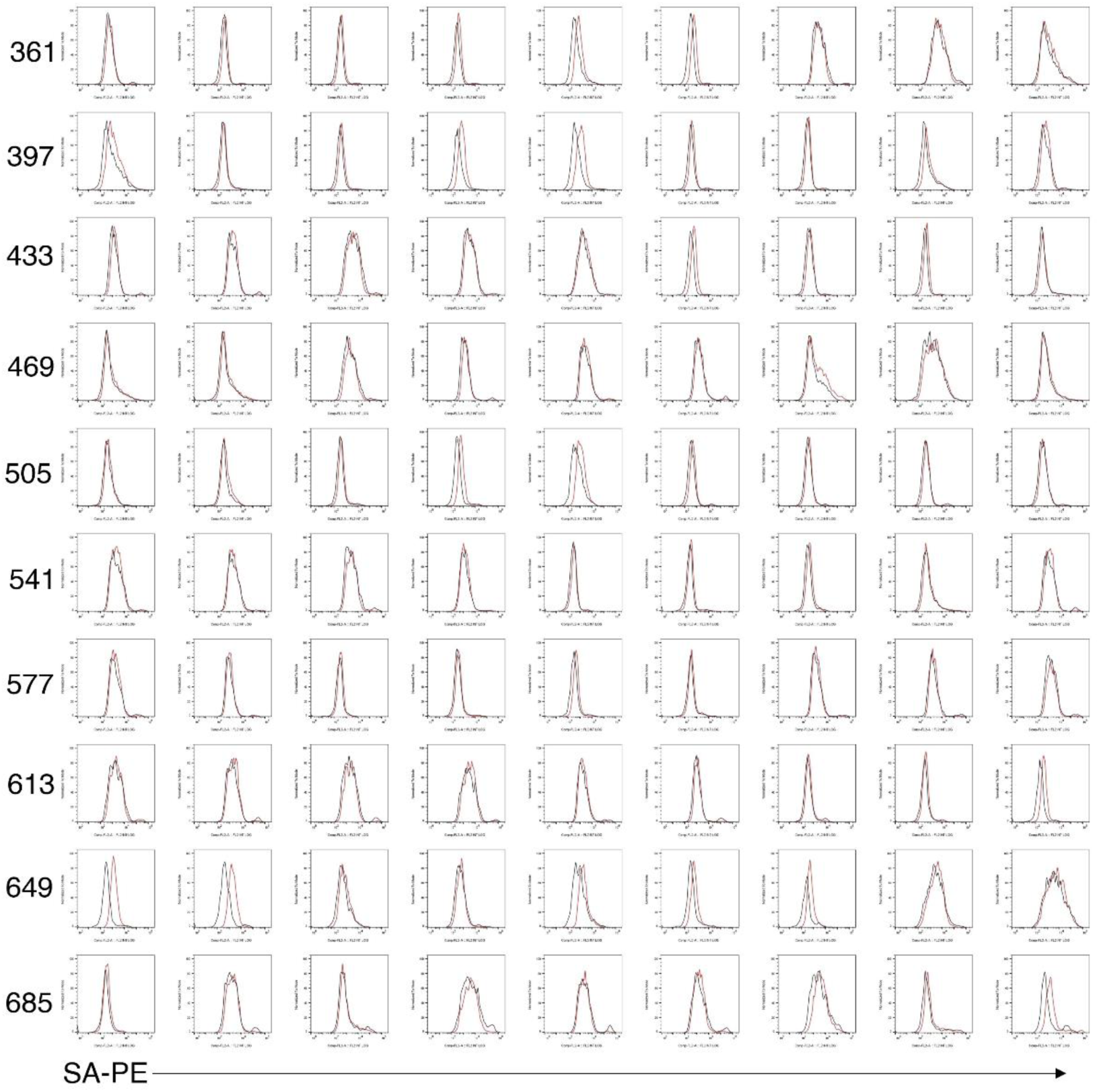
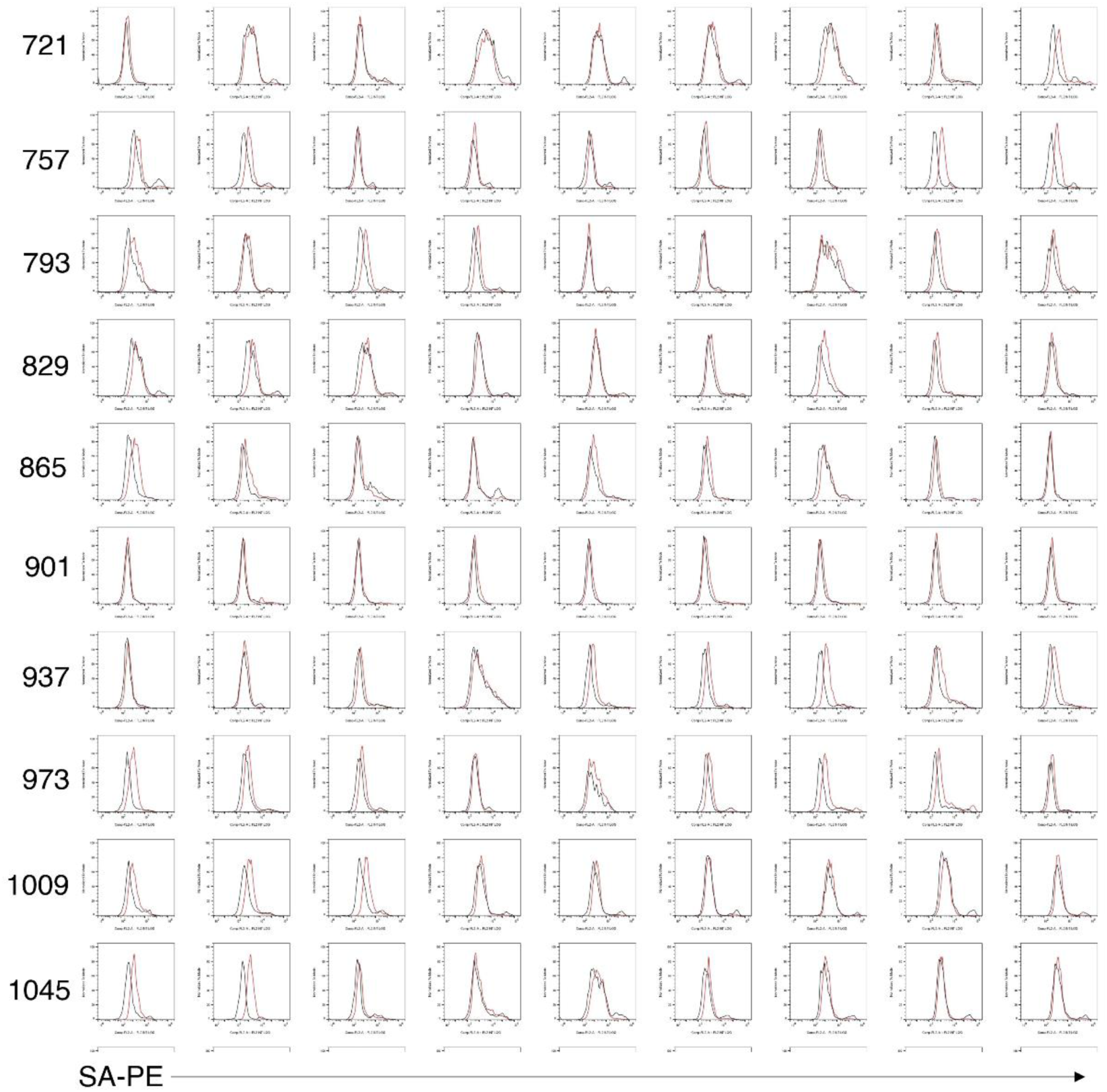
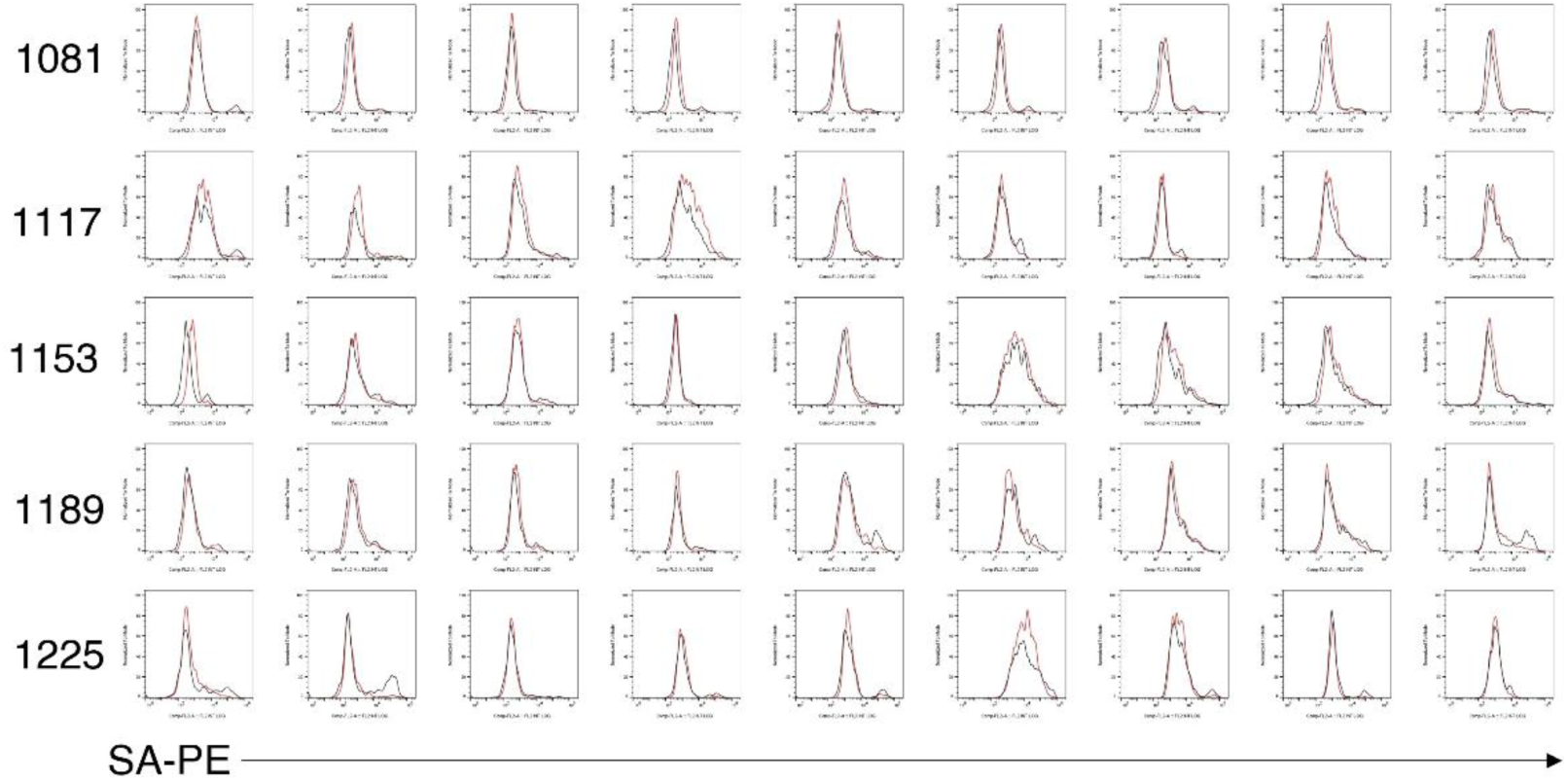
Representative flow cytometry data Representative histogram of all peptides binding to HLA-DRB1*15:01. The black line is HLA null cells. The red line is DRB1*15:01-expressing cells.

**Supplemental Figure 4.**
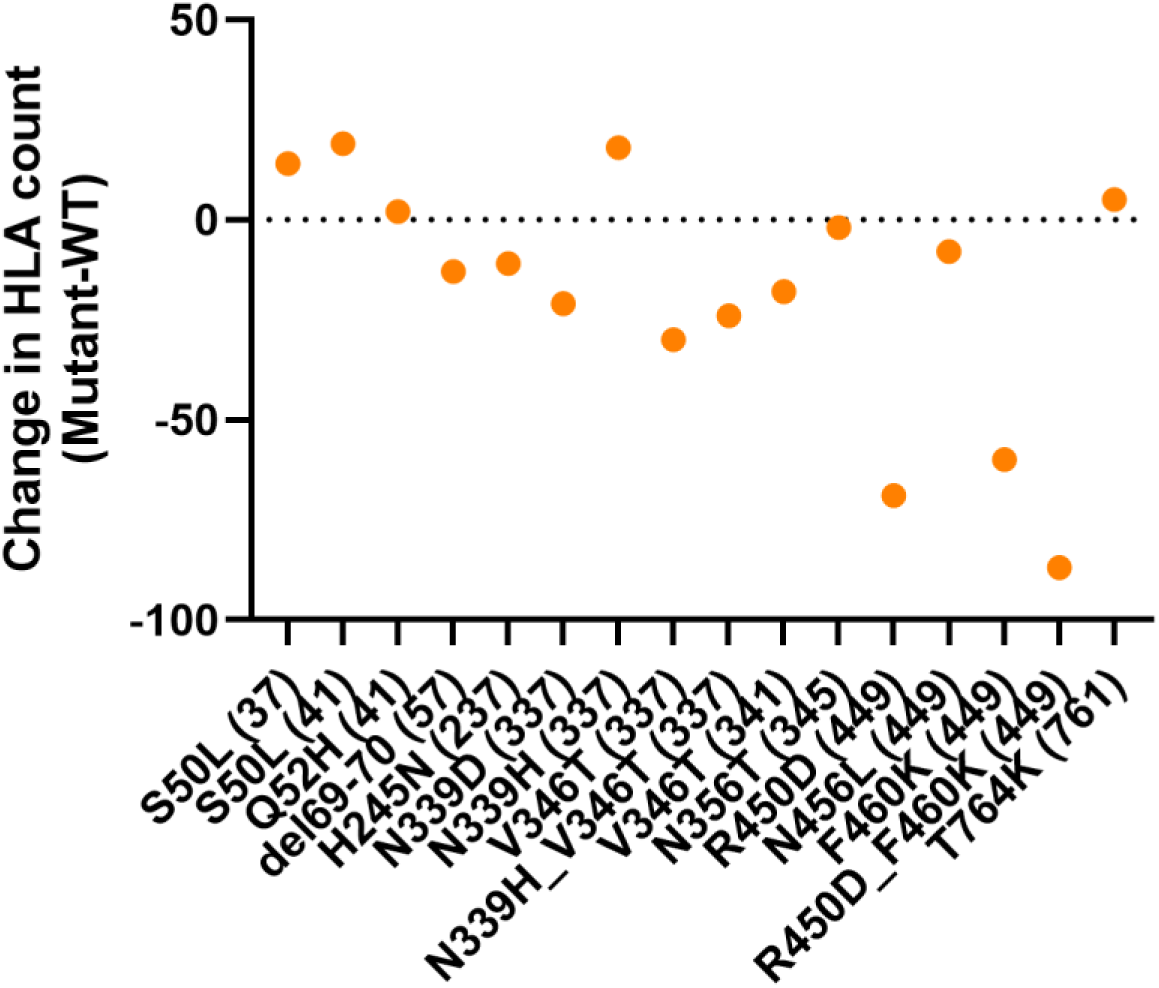
Effects of mutations on predicted peptide binding. The number of HLAs which are predicted by any method to bind the mutant peptide was subtracted from the number of HLAs which are predicted by any method to bind the WT peptide was determined. The numbers in parenthesis denote the peptide containing the mutation.

## Acknowledgements

We would like to Satoru Iwai for helping produce the HLA-expressing mouse hybridoma cells. We would also thank Carla Günther for productive discussions. This work was supported by the Japan Agency for Medical Research and Development (JP223fa627002, JP223fa727001, JP23ym0126049 (SY)), and Japan Society for the Promotion of Science Grants-in-Aid for Scientific Research (JP20H00505, JP22H05182, JP22H05183 (SY)).

## Author Contributions

Charles Schutt, conceptualization, investigation and manuscript preparation, Deren Birol, investigation, Xiuyuan Lu, conceptualization and manuscript editing, Sho Yamasaki, conceptualization, supervision, resources and funding acquisition.

